# High-parameter phenotypic characterization reveals a subset of human Th17 cells that preferentially produce IL17 against *M. tuberculosis* antigen

**DOI:** 10.1101/2023.01.06.523027

**Authors:** Paul Ogongo, Anthony Tran, Florence Marzan, David Gingrich, Melissa Krone, Francesca Aweeka, Cecilia S Lindestam Arlehamn, Jeffrey N. Martin, Steven G. Deeks, Peter W. Hunt, Joel D. Ernst

**Affiliations:** Division of Experimental Medicine, University of California, San Francisco, CA, USA; Department of Tropical and Infectious Diseases, Institute of Primate Research, Nairobi, Kenya; Drug Research Unit, Department of Clinical Pharmacy, School of Pharmacy, University of California, San Francisco, CA, USA; Department of Epidemiology and Biostatistics, University of California, San Francisco, CA, USA; Center for Infectious Disease and Vaccine Research, La Jolla Institute for Immunology, La Jolla, CA 92037, USA; Division of HIV, Infectious Diseases, and Global Medicine, University of California, San Francisco, CA, USA

**Keywords:** Interleukin-17, CD4 T-cells, antigen-responsive, immunity, Tuberculosis, ART-suppressed, HIV, kynurenine pathway

## Abstract

**Background:** Interleukin 17 producing CD4 T cells contribute to the control of *Mycobacterium tuberculosis (Mtb)* infection in humans; whether infection with Human Immunodeficiency Virus (HIV) disproportionately affects distinct Th17 cell subsets that respond to *Mtb* is incompletely defined.

**Methods:** We performed high-definition characterization of circulating *Mtb*-specific Th17 cells by spectral flow cytometry in people with latent TB and treated HIV (HIV-ART). We also measured kynurenine pathway activity by LC/MS on plasma and tested the hypothesis that tryptophan catabolism influences Th17 cell frequencies in this context.

**Results:** We identified two subsets of Th17 cells: subset 1 defined as CD4^+^Vα7.2^-^CD161^+^CD26^+^ and subset 2 defined as CD4^+^Vα7.2^-^CCR6^+^CXCR3^-^ cells of which subset 1 was significantly reduced in LTBI with HIV-ART, yet *Mtb*-responsive IL17-producing CD4 T cells were preserved; we found that IL17-producing CD4 T cells dominate the response to *Mtb* antigen but not CMV antigen or staphylococcal enterotoxin B (SEB); and tryptophan catabolism negatively correlates with both subset 1 and subset 2 Th17 cell frequencies.

**Conclusions:** We found differential effects of ART-suppressed HIV on distinct subsets of Th17 cells, that IL17-producing CD4 T cells dominate responses to *Mtb* but not CMV antigen or SEB, and that kynurenine pathway activity is associated with decreases of circulating Th17 cells that may contribute to tuberculosis immunity.

## 1 Introduction

Increasing evidence indicates that CD4 T cells that produce interleukin 17 (IL17), broadly termed T helper 17 (Th17) cells, contribute to the control of TB. In *Mtb*-infected adolescents, IL17 transcriptional signatures decreased in the blood of those who progressed to active TB compared to non-progressors [1]; another study found a subset of CD4 T cells that produce IL17 in response to *Mtb* antigens that is less abundant in TB progressors than in non-progressors [2]. *Mtb*-responsive CD4 T cells producing IL17 are enriched in human lungs compared to matched blood and inversely correlated with plasma IL-1β, suggesting a role in the control of *Mtb* [3]. IL17 producing CD4 T cells are depleted in HIV infection and their depletion contributes to the progression to AIDS through breakdown in intestinal mucosal barrier function reviewed in [4,5], in mechanisms that depend in part on the expression of CCR5 by IL17 producing CD4 T cells.

Several approaches are used to identify T helper 17 cells (herein denoted ‘Th17’ cells to include all subsets regardless of criteria used). IL17 production upon stimulation is the canonical signature of Th17 cells [6,7]. Although this identifies Th17 cells, it does not classify all cells in the Th17 lineage, and other cells can produce IL17 [8–11]. In the blood of healthy donors and synovial fluid of rheumatoid arthritis patients, a CCR6^+^ subset contained all IL17-producing T cells expressing *RORC* mRNA [12], the transcription factor that supports Th17 differentiation in humans. The use of CCR6 as a marker of Th17 cells has been replicated in naïve cord blood [13], inflammatory diseases [14], and infections like TB [15]. The cell surface markers CD26 [16–19] and CD161 [20–23] have also been used to identify Th17 cells. CD26^hi^ CD4^+^ T cells were demonstrated to co-express CD161 and CCR6 and to be enriched for the production of IL17 [16]. In transcriptomic studies, Th17 cells are identified by expression of *RORC*, *IL23R* (supports the expansion of IL17-producing cells) and *IL17* mRNA.

Host metabolism impacts immune responses [24] including by driving cell differentiation. One example is the kynurenine pathway of tryptophan catabolism whose products, collectively called kynurenines, inhibit Th17 while promoting Treg development [25–28]. In TB, circulating tryptophan (Trp) concentrations decline in persons with LTBI progressing to active TB [29] and Trp concentration increases with TB treatment [25]. Similarly, HIV infected persons have low circulating Trp concentration with a link to pathogenesis [30]. Plasma kynurenine/tryptophan ratios (K/T) can distinguish humans with active TB from those without and are significantly higher in MDR-TB patients with or without HIV coinfection compared to controls [25]. Plasma K/T is also elevated in people living with HIV (PWH) compared to controls, especially in those who have progressed to AIDS [27,31]. While a link between K/T and Treg/Th17 ratio has been established in HIV, similar data are lacking in TB.

Here, we characterized subsets of Th17 cells defined by distinct criteria in participants with ART-suppressed HIV (HIV-ART), to determine whether IL17-producing CD4 T cells that respond to *Mtb* antigens are altered in HIV-ART, and to test the hypothesis that indoleamine 2,3-dioxygenase (IDO) activity and generation of kynurenines are associated with Th17 differentiation in HIV-ART.

## Materials and Methods

### Participants and sample collection

PBMC and plasma were obtained from adults with or without HIV enrolled in the UCSF SCOPE cohort [27,32]. The participants selected for the current study were tested for sensitization to *Mtb* by tuberculin skin test (TST). All participants gave written informed consent using protocols approved by the UCSF Institutional Review Board. Blood was collected into EDTA-containing tubes for cell counts and into ACD-containing tubes for purification of plasma and PBMCs. Processed plasma was stored at -80°C while PBMCs were cryopreserved in liquid nitrogen. Demographic characteristics of the participants are in Tables 1 and 2. We did not have samples from all the participants in the longitudinal study (cohort 2/table 2) at the start of isoniazid (INH) therapy.

**Table 1:** Cohort 1 – cross sectional -(Characteristics of participants latent TB and with or without HIV)\.

|  |  |  |  |
| --- | --- | --- | --- |
| **Characteristic** | **HIV negative (n =7)** | **HIV positive (n = 21)** | **P** |
| Age, median (IQR), y | 58 (44 -61) | 47 (38 - 58) | 0.5395 |
| Sex, Male (%) | 7 (100) | 21 (100) |  |
| Race/Ethnicity (n, (%)) |  |  |  |
| Black | 6 (86) | 8 (38) |  |
| Asian | 1 (14) | 2 (10) |  |
| White | 0 | 7 (30) |  |
| Middle East | 0 | 1 (5) |  |
| Multiracial | 0 | 2 (10) |  |
| Pacific Islander | 0 | 1 (5) |  |
| Combined antiretroviral therapy (Yes/No) | No | Yes |  |
| CD4 Counts, median (IQR), cells/mm3 | 1297 (707 -1349) | 640 (487 - 906) | 0.0082 |
| Plasma viral load (copies/mL) | None | < 40 |  |

**Table 2:**
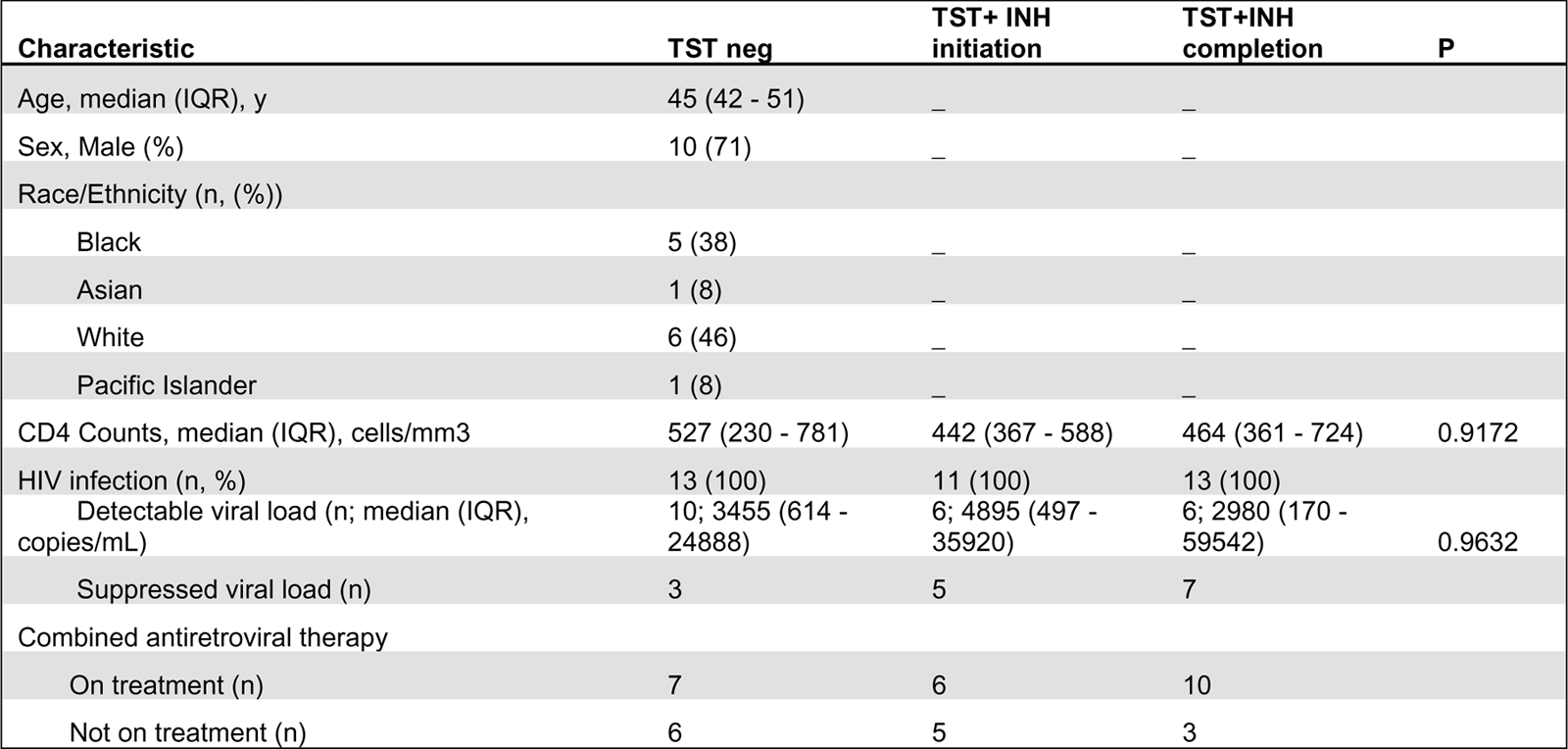
Cohort 2 – longitudinal - (Characteristics of participants with HIV before and after LTBI diagnosis and treatment)

### Plasma Tryptophan and Kynurenine measurement

Tryptophan and Kynurenine concentrations were measured by liquid chromatography-mass spectrometry as previously described [27,31].

### PBMC antigen stimulation and Intracellular cytokine staining

Cryopreserved PBMCs were thawed in a 37°C water bath and transferred into pre-warmed R10 media (RPMI 1640 containing L-glutamine with 10% FBS, 1%PenStrep and 1% Hepes). Cells were centrifuged at 900 xg for 5 minutes at room temperature, supernatants discarded, and cells resuspended before adding 5ml of warm R10 and transferred to a 6-well culture plate and resting overnight in 37°C/5%CO_2_ incubator. After resting, 1x10^∧6^ live cells resuspended in 200µL R10 were transferred to each well of a 96-well round bottom plate, and duplicate wells were stimulated with *Mtb* peptide mega pool (Mtb300) [33](2µg/ml), CMV pp65 peptide (1µg/ml) or Staphyloccal enterotoxin B (SEB) positive control (1µg/ml) in the presence of costimulating antibodies anti CD28 (1µg/ml) and anti CD49d (1µg/ml) (BD). *Mtb* peptide megapool represents commonly recognized epitopes from 90 *Mtb* antigens restricted by HLA class II-restricted CD4 T cells in diverse populations and across species [3,34–42]. CMV pp65 (PepTivator® CMV pp65 – Milteny Biotec) is a peptide pool covering the complete sequence of the pp65 protein of cytomegalovirus. Negative controls received no stimulation. After two hours, GolgiStop and GolgiPlug (BD Biosciences) were added to one well of each of the stimulation conditions, and the cells were incubated for an additional 18 hours.

After 20-hour total stimulation, cells were washed, stained with Live/Dead Fixable Blue Dead Cell stain kit (Invitrogen), washed again, then surface antibody cocktail (αCD3 BV510 clone UCHT1, 1:80; αCD8 BV570 clone RPA-T8, 1:80; αCCR7 BV785 clone G043H7, 1:20; αCD95 Alexa Fluor700 clone DX2, 1:80; αCCR6 FITC clone G034E3, 1:80; αCXCR3 BV 605 clone G025H7, 1:40; αCD161 APC-Fire750 clone HP-3G10, 1:80; αCD69 BV650 clone FN50 ,1:20; αCD137 BV711 clone 4B4-1, 1:20, αOX40 PE-Cyanine7 clone Ber-ACT35, 1:40 and αTRAV1-2 (Vα7.2) BV421 clone 3C10, 1:40 (all from BioLegend), αCD4 BUV496 clone SK3, 1:80, αCD45RA BUV395 clone 5H9, 1:40, αCD27 BUV615 clone L128, 1:80, αCD25 BUV563 clone 2A3, 1:80, αCD39 BUV737 clone TU66, 1:80 and αCD26 BUV805 clone M-A261, 1:80 (all BD Biosciences)) diluted in Brilliant Violet buffer (BD) was added then kept in the dark for 20 minutes at room temperature., After washing, cells were fixed and permeabilized using eBioscience FOXP3/Transcription Factor staining kit (Invitrogen). Cells were washed twice with 1X eBioscience Perm diluent, then resuspended in intracellular antibody mix (αRORγT Alexa Fluor 647 Clone Q21-559, 1:40, αIFNγ BB700 clone 2B7, 1:160 (both from BD Biosciences), αIL17 PE clone BL168, 1:20, αT-bet PE-Dazzle 594 clone 4B10, 1:40 (both from BioLegend) and αFoxP3 PE-Cyanine5.5 clone PCH101, 1:40 (Thermofisher)) in eBiosciences perm diluent. Cells were then washed (centrifuged at 900 xg for 5 minutes at room temperature) twice with 1X eBioscience Perm diluent and fixed in 2% PFA. Data was acquired on a 5-Laser Aurora Spectral Flow cytometer (Cytek).

### Data and statistical analysis

Spectral flow fcs files were initially analyzed using SpectroFlo v3.0 (Cytek) for unmixing and autofluorescence correction. Identification of T cell subsets from unmixed fcs files was performed using Flowjo v10 (Flowjo LLC). Antigen-specific cytokine levels are reported after subtraction of values of unstimulated cells for each participant. K/T ratio was determined from the concentrations of tryptophan and kynurenines after quality control checks. Non-parametric tests were used for comparison between two groups for paired (Wilcoxon test) or unpaired (Mann-Whitney test) and p < 0.05 was considered significant. Spearman correlation was used to test the association between parameters with a p < 0.05 considered significant. All statistical analyses used Prism 9.0 (GraphPad).

## Results

### Subsets of circulating T helper 17 cells are disproportionately reduced in LTBI with treated HIV

To study the effects of LTBI and ART-treated HIV (HIV-ART) on Th17 cells defined by different criteria, we designed a spectral flow cytometry panel encompassing cytokines, chemokine receptors, transcription factors, and cell surface markers (Figure S1 and Figure 1A&C). We used the following nomenclature: (a) subset 1 is CD26^+^CD161^+^ [16,17]; (b) subset 2 is CCR6^+^CXCR3^-^ [12,15]; and (c) Th1* (T1T17) is CCR6^+^ CXCR3^+^ [15,43]. We first gated out Vα7.2^+^ CD4^+^ T cells to exclude MAIT cells that share certain markers with Th17 cells [44–49], and since MAITs can produce IL17 [50].

**Figure 1.**
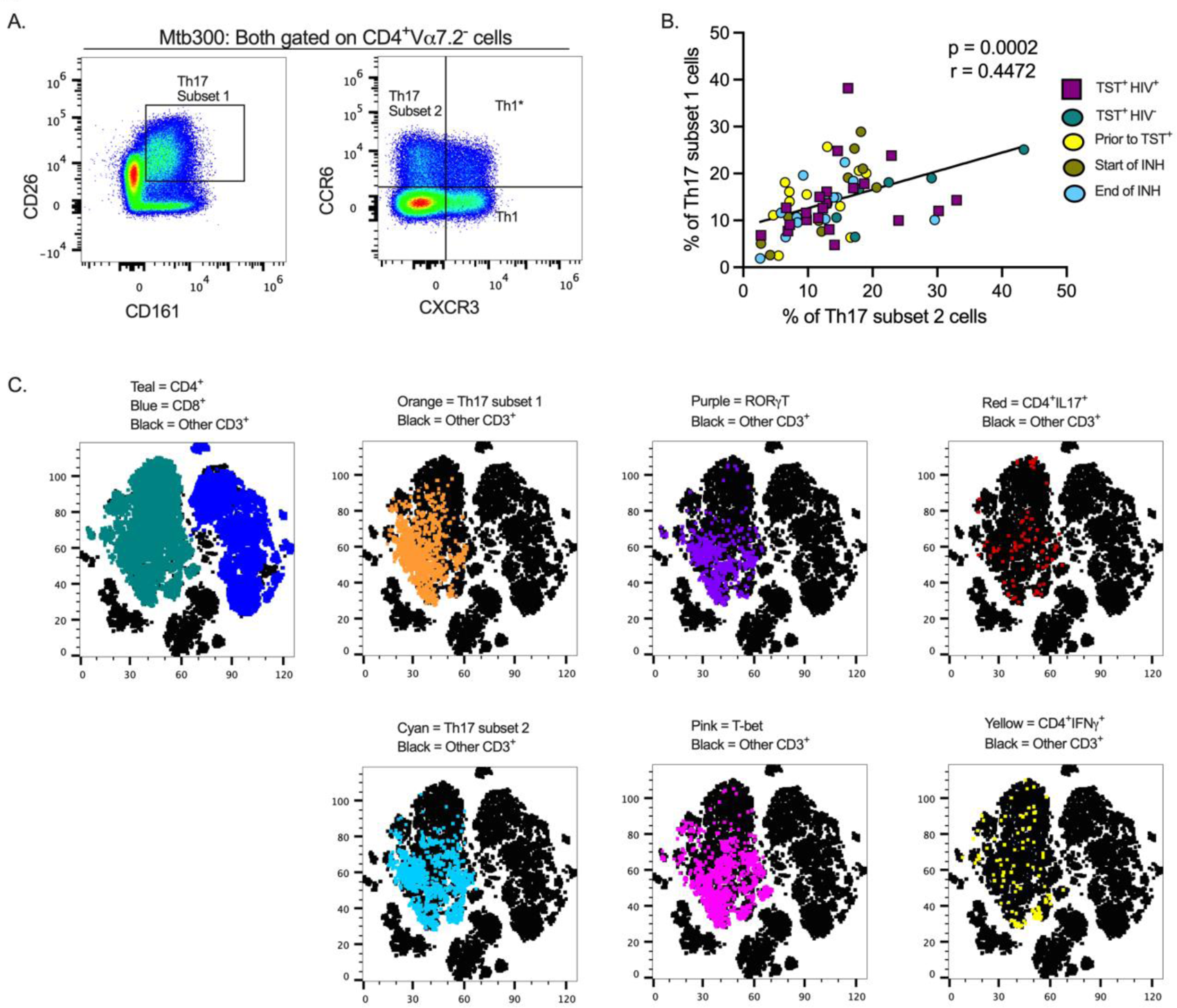
Different definitions of T helper 17 cells reveal high but incomplete concordance. **(A)** Representative flow cytometry plots and gating schemes for definition of Th17 cells by published marker criteria independent of cytokine expression. Left panel, chemokine receptor CCR6 and CXCR3 expression used to distinguish subset 2, Th1*, and Th1 non-MAIT CD4 T cells. Right panel: Use of surface expression of CD26 and CD161 to identify subset **1** non-MAIT CD4 T cells. Cells in both panels were stimulated with the MTB300 antigenic peptide pool. **(B)** High correlation of subset 1 (CD2s+co161+) and subset 2 (CCR6+CXCR3-) non-MAIT CD4 T cells in cell populations from participants with varying TB and HIV status. Spearman correlation p and r values are shown for results pooled from all participants. (C) t­stochastic neighbor embedding (t-SNE) analysis reveals incomplete concordance of transcription factor (RORyt and T-bet) and cytokine (IFNy and IL17) production and surface phenotypes of non-MAIT co4+ T cells. In this analysis, protein transport inhibitors (golgi stop and golgi plug) were not added to PBMC during antigen stimulation thus cytokine detection was not optimal. Prior to Tsr = participants were considered Mtb unexposed due to a negative TST test; INH = isoniazid therapy. Statistic: Spearman correlation.

We also stained for RORγT, the Th17 lineage defining transcription factor [51], and used unsupervised clustering and t-distributed stochastic neighbor embedding (tSNE) to determine whether subset 1 and subset 2 Th17 cells express RORγT and/or produce IL17 upon stimulation with an *Mtb* peptide mega pool (Mtb300) (Figure 1C). To identify Th1* (also termed T1T17) cells that respond to *Mtb* antigens [15], we included T-bet, the Th1 lineage transcription factor [52]. We found a positive and significant association (r = 0.4472, p = 0.0002) (Figure 1B) between subset 1 and subset 2 Th17 cells, and found that cells in both populations express RORγT and produce IL17 (Figure 1C). Therefore, subset 1 and subset 2 Th17 cells express other properties of bona fide Th17 cell populations. T-bet expressing cells were present within subset 1 and subset 2 Th17 cell populations suggesting a Th1* population that expresses both CCR6 and CXCR3 and likely produce both IFNγ and IL17 (Figure 1C). Thus, we identified the Th17 subsets based on chemokine receptors (CCR6^+^ in the absence of CXCR3 expression) or surface markers (CD26 and CD161) co-expression.

As expected, circulating CD4 T cells were reduced in HIV-ART compared with HIV-uninfected participants, (Figure 2A). We first phenotyped CD4^+^ T cell populations based on CD45RA and CCR7 [3] to identify naïve, central memory, effector memory, and terminally differentiated T cells (Figure S2 A, left). The predominant populations were naïve and central memory T cells and there was no difference in the frequency of cells in these memory populations in participants with or without HIV (Figure 2B). CD4^+^CD45RA^+^CCR7^+^ (naïve T cells) are a heterogeneous population, including cells that produce cytokines upon short-term stimulation [53], so we characterized the naïve T cells further based on co-expression of CD95 and CD27 to identify cells with a stem cell-like memory phenotype (Figure S2 A, right). We found no difference in frequencies of CD95^+^CD27^+^ stem cell-like memory cells or CD95^-^CD27^+^ ‘true naïve’ cells in those with or without HIV (Figure S2 B). Thus, whereas individuals with treated HIV have lower overall frequencies of circulating CD4^+^ T cells than those without HIV (Tables 1 and 2, Figure 2A), their memory states do not differ. These observations are in accord with studies that antiretroviral therapy induces a burst in CD4 T cell memory regeneration and general reconstitution of CD4 T cell count relative to pre-treatment time points [54–56].

**Figure 2.**
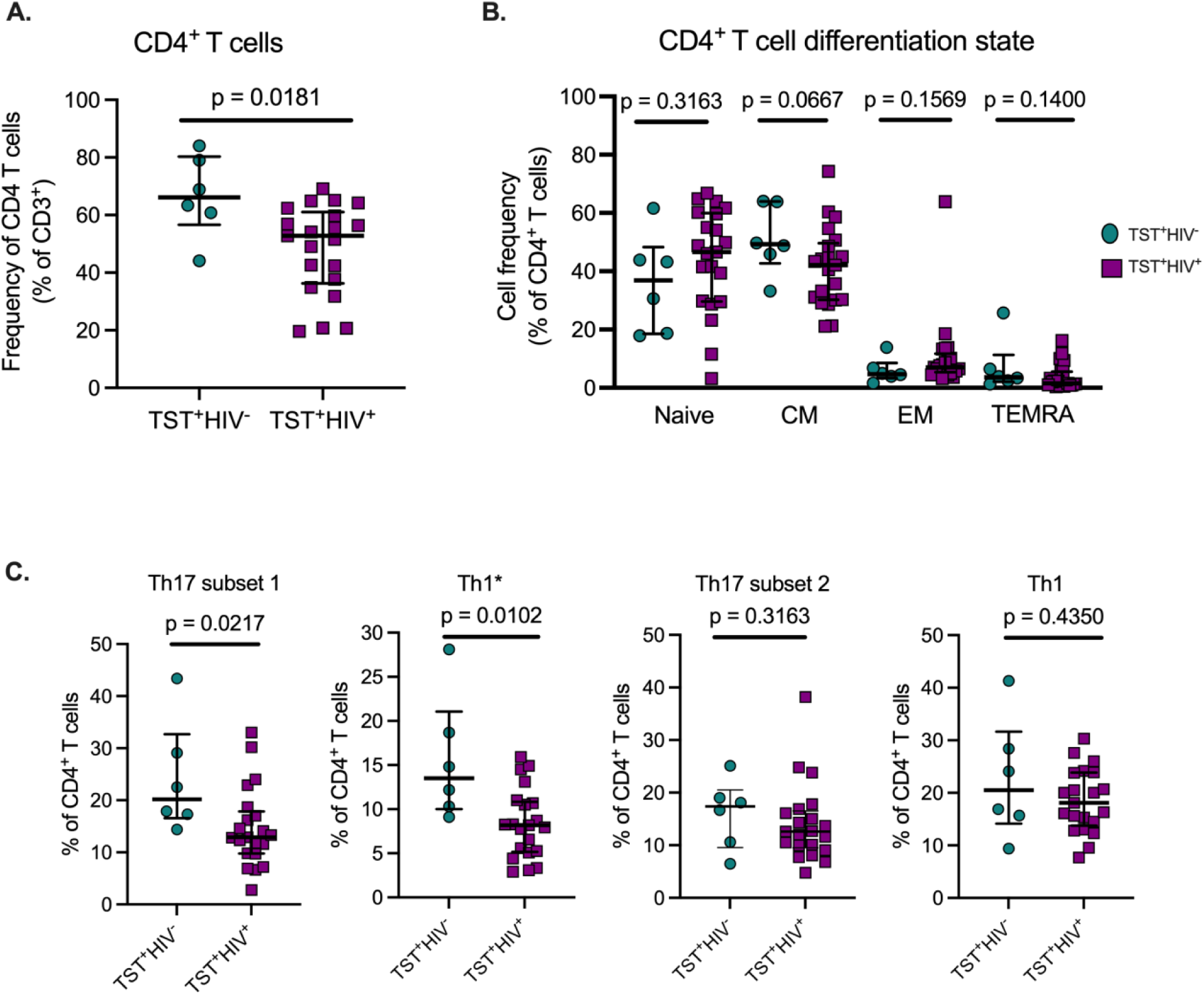
ART treated HIV infection is associated with disproportionate depletion of distinct subsets of T helper cells. **(A)** Reduced frequency of bulk CD4 T cells in HIV+ vs HIV· participants in Cohort 1. **(B)** ART-treated HIV is not associated with differential distribution of CD4 T cells in memory T cell subsets. Naive: CD45RNCCR7+; Central Memory (CM): CD45RACCRr); Effector Memory: CD45RACCR7·; TEMRA: CD45RA+ccR?·). **(C)** Disproportionate reduction in frequencies of subset 1 and Th1* but not subset 2 or Th1 cells in ART-treated HIV. Subset 1 = co4+va?.2·co26+co161+; Th1* (also termed T1T17): co4+va?.2·ccR6+cxcR3+; subset 2: co4+va?.2·ccR6+cxcR3·; and Th1: co4+va1.2-ccR6·cxcR3+. Statistical comparisons were by Mann-Whitney test.

Next, we quantitated Th17 cells in those with LTBI without HIV or with HIV-ART and found significantly lower frequencies of subset 1 (CD4^+^Vα7.2^-^CD26^+^CD161^+^) and Th1* (CD4^+^Vα7.2^-^ CCR6^+^CXCR3^+^) cells (expressed as % of CD4^+^ T cells) in HIV-ART than in those without HIV (Figure 2C). Although subset 2 cells (CD4^+^Vα7.2^-^CCR6^+^CXCR3^-^) and Th1 (CD4^+^Vα7.2^-^CCR6^-^CXCR3^+^) cells were also present at lower frequencies in HIV-ART, the difference was not significant (Figure 2C). These data demonstrate that CD26^+^CD161^+^ (subset 1) circulating Th17 cells are disproportionately depleted in HIV-ART, while CCR6^+^ (subset 2) Th17 cells and Th1* (T1T17) cells are relatively preserved.

Th17 and regulatory T cells (Tregs) develop under related but distinct cytokine environments [57]. Tregs contribute to immune homeostasis by maintaining unresponsiveness to self-antigens, suppressing exaggerated immune responses, and promoting epithelial tissue integrity [58]. Human Tregs are identified by the expression of high levels of CD25 and the transcription factor FoxP3 [59]. Since the kynurenine pathway has differential effects on Th17 and Treg differentiation, we characterized Treg cells (Figure S3A, left) and found that the frequency of circulating Treg was similar in HIV-ART and those without HIV (Figure S3B). We also found that the frequency of activated (CD39^+^) Tregs did not differ in the two groups (Figure S3A, right) although there was considerable variation in CD39^+^ Treg, especially in HIV-ART (Figure S3 C). We also compared the ratio of Th17 subset 1 and T17 cells to Tregs between the groups and found a trend toward a lower Th17 subset1/Treg in HIV-ART than in those without HIV (Figure S3D), but the difference was not significant, and T17/Treg ratios were similar in HIV-ART and those without HIV (Figure S3E). Taken together, we found that certain CD26^+^CD161^+^ Th17 cell subset is disproportionately reduced in LTBI with treated HIV, suggesting that even effective treatment of HIV does not reconstitute all Th17 cell subsets equally. We did not find differences in Treg by HIV status.

### Mtb antigen-responsive IL17-producing cells are enriched in subset 1 compared with subset 2 Th17 cell populations in HIV-ART

Having established that subset 1 (CD26^+^CD161^+^) and subset 2 (CCR6^+^) Th17 cells are bona fide Th17 populations, we determined whether subset 1 and subset 2 cells differ in IL17 production after *Mtb* antigen stimulation. HIV-ART participants with LTBI had significantly more IL17^+^ cells in subset 1 Th17 cells than in subset 2 Th17 cells upon stimulation with the Mtb300 peptide pool (Figure 3A, left). All except one participant had IL17^+^ cells in subset 1 cells compared with 11 of 21 participants with IL17^+^ cells in subset 2 Th17 cells (Figure 3A, left). We did not have enough cells from participants without HIV (cohort 1) for cytokine analysis. We confirmed this observation in cohort 2; regardless of the sampling time point, there were significantly more IL17^+^ cells in subset 1 Th17 cells than in subset 2 Th17 cells (Figure 3A, right). We performed a similar analysis after stimulation with SEB and found an opposite pattern: IL17 production was significantly higher in subset 2 than in subset 1 Th17 cells (Figure 3B). Additionally, CMV peptide stimulation induced equivalent production of IL17 by subset 1 and subset 2 Th17 cells, although there was a trend towards higher responses in subset 2 Th17 cells (Figure S4A). These data reveal enrichment of IL17 production to *Mtb* antigens by a subset of Th17 cells marked by expression of CD26 and CD161 (subset 1) which are distinct from IL17-producing cells that are marked by expression of chemokine receptor CCR6 (subset 2) and those that produce IL17 in response to SEB or CMV.

**Figure 3.**
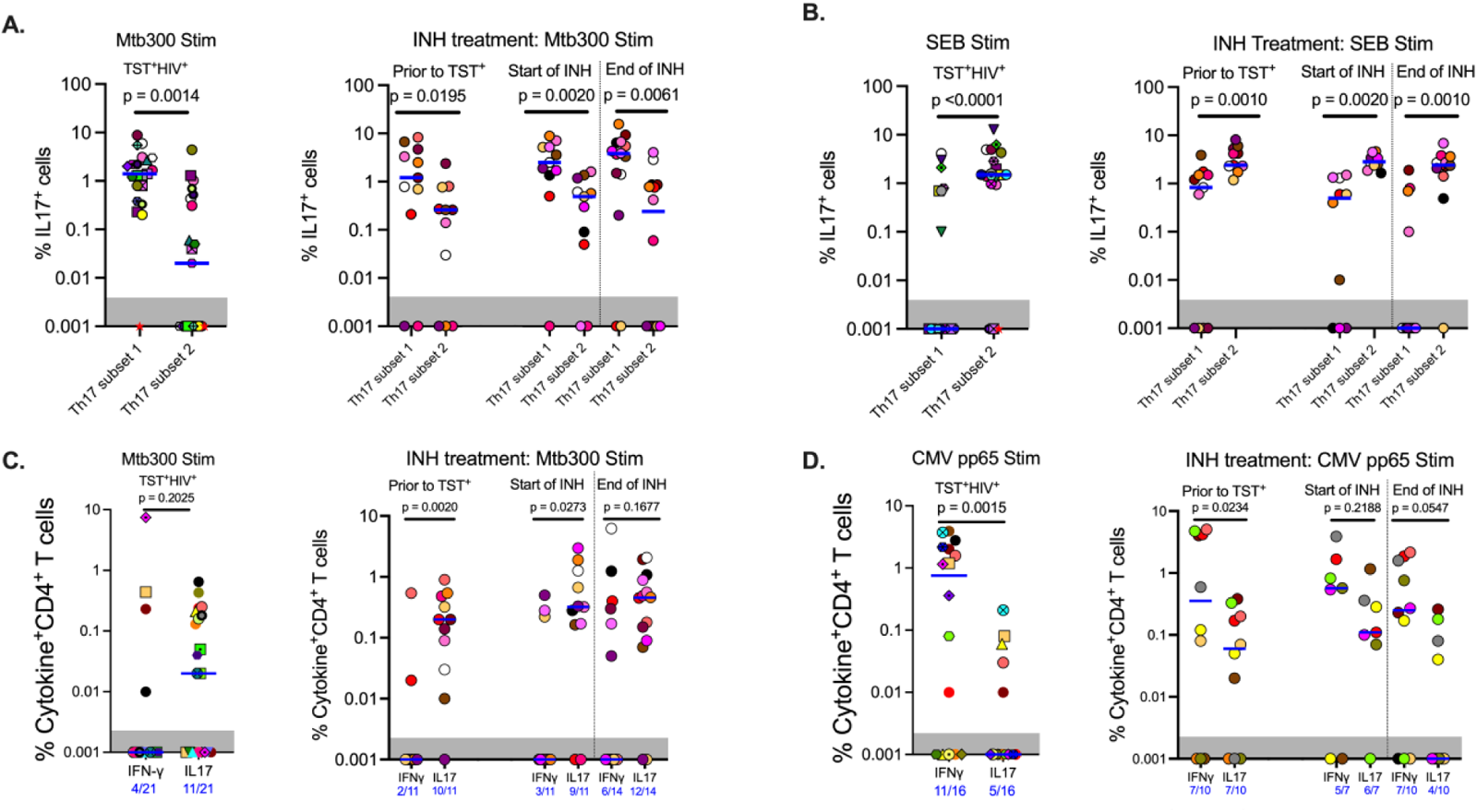
In contrast to responses to other stimuli, *Mtb* antigen responsive IL17 expression is enriched in subset 1 (CD26^+^CD161^+^) rather than subset2 (CCR6^+^CXCR3) Th17 cells. **(A)** Left panel (Cohort 1): CD4 T cells that produce IL17 in response to MTB300 are predominantly in subset 1. Right panel (Cohort 2): MTB300-stimulated IL17 production predominates in subset 1 rather than subset 2 cells regardless of TST status and is not altered by INH preventive therapy. **(B)** Left panel (Cohort 1): In contrast to MTB300-responsive cells, IL17 production predominates in subset 2 cells in response to SEB activation. Right panel (Cohort 2): INH preventive therapy does not alter the pattern of IL17 production by SEB-stimulated subset 2 or subset 1 Th17 cells. **(C)** Left panel (Cohort 1): In ART-treated HIV^+^ participants, CD4 T cells that produce IFNγ in response to MTB300 stimulation are present at low frequencies. Right panel (Cohort 2): **(D)** Left panel (Cohort 1): CMV pp56 antigen stimulation induces higher frequencies of IFNγ than IL17-producing CD4 T cells. Right panel (Cohort 2): INH treatment does not alter CD4 T cell responses to CMV pp65 antigen stimulation. Statistics: Wilcoxon test.

### Mtb antigen-responsive IL17-producing CD4^+^ T cells are relatively preserved in people with LTBI and HIV-ART

Next, we assessed IFNγ and IL17 production after Mtb300 stimulation. In HIV-ART, only 4 of 21 (19%) had detectable IFNγ responses compared with 11 of 21 (52%) who had detectable IL17 responses (p = 0.0516, Fisher’s exact test). Additionally, the median IL17 response was higher than the median IFNγ response although this did not reach significance (Figure 3C, left). Similar IL17 responses to *Mtb* antigen stimulation were observed in people with HIV before and after LTBI diagnosis and treatment. Treatment of LTBI did not change either IFNγ or IL17 responses although the response magnitudes varied by participant (Figure 3C, right). Similarly, more participants had detectable IL17 responses than detectable IFNγ responses at all 3 sampling times: prior to being TST positive, and at INH initiation and treatment completion: 2/11 vs 10/11, 3/11 vs 9/11 and 6/14 vs 12/14, respectively (Figure 3C, right).

To determine whether the predominant production of IL17 over IFNγ was specific to *Mtb* antigens, we used a CMV peptide pool to stimulate PBMC from participants (Tables 1 and 2) who had sufficient cells (Figure 3D). This revealed a pattern distinct from that with Mtb300: participants with LTBI and HIV-ART (cohort 1) exhibited higher magnitude and higher prevalence of IFNγ than IL17 responses (IFNγ: 11/16 (69%) vs IL17: 5/16 (31%)) (Figure 3D, left) (p = 0.0756, Fisher’s exact test). Similarly, in Cohort 2 there was a higher magnitude of IFNγ responses than IL17 responses at all time points, reaching statistical significance at baseline (prior to TST^+^) timepoint (Figure 3D – right). As with CMV and in contrast to Mtb300, SEB induced higher magnitude IFNγ than IL17 responses (Figure S4B). Together, these data demonstrate that in LTBI and HIV-ART, IL17 responses to *Mtb* are relatively preserved compared with IFNγ responses, and this is distinct compared to CMV and SEB responses. This is also contrary to the commonly held impression that IFNγ production is the predominant response to *Mtb*.

### IDO activity is increased in people with LTBI and HIV-ART, and negatively correlates with circulating Th17 cells

We quantitated tryptophan and kynurenine concentrations in plasma of participants with LTBI with or without HIV-ART and inferred IDO activity based on the K/T ratio. We found significantly lower plasma tryptophan concentrations in participants with HIV-ART compared with those without HIV (Figure 4A) and kynurenine concentrations were higher in HIV-ART but did not achieve statistical significance when compared with the HIV uninfected group (Figure 4B). Finally, there was a significantly higher K/T ratio in people with HIV-ART compared to those without HIV (Figure 4C). Although we cannot ascertain the contribution of LTBI to the observed IDO activity because we did not have participants who were not exposed to *Mtb*, these observations agree with prior reports showing that IDO-1 mediated tryptophan catabolism is elevated in PWH, including those initiated on early treatment and ART-suppressed viral loads [31].

**Figure 4.**
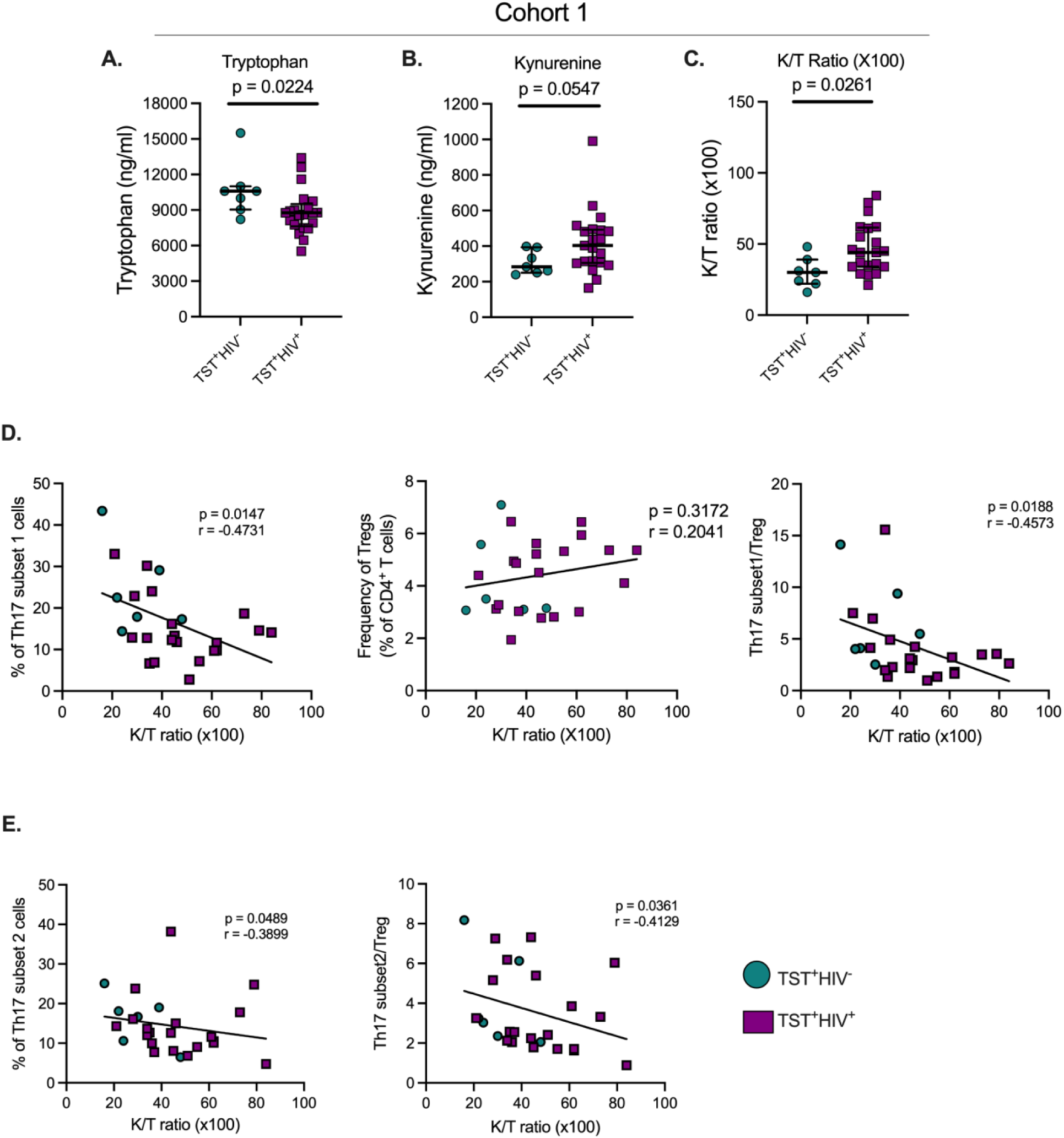
In TS-r+ participants, HIV infection is associated with increased indoleamine-2,3-dioxygenase (IDO) activity, which negatively correlates with Th17 cell subsets. **(A)** Plasma tryptophan concentrations; **(B)** Plasma kynurenine concentrations; **(C)** Plasma Kynurenine (K)/Tryptophan (T) ratios as a reflection of ID0 activity. **(D)** Left panel: correlation of subset 1 Th17 cell frequencies with plasma KIT ratios; Middle panel: Correlation of regulatory T cells and plasma KIT ratios; Right panel: Correlation of the ratio of subset 1/Treg ratios and plasma KIT ratios. **(E)** Left panel: correlation of subset 2 Th17 cell frequencies with plasma KIT ratio; Right panel: Correlation of the ratio of subset 2/Treg ratios and plasma KIT ratios. Statistical analyses were by Mann-Whitney (panels A-C) or Spearman correlation (D and E).

Since kynurenines favor the development of Tregs while suppressing Th17 generation [27], we determined the correlation of plasma K/T and Th17 cells, Tregs, and Th17/Treg ratios in Cohort 1 participants. This revealed a significant inverse relationship between K/T and circulating subset 1 and subset 2 (Figure 4D & E), while there was no correlation between K/T and Tregs (Figure 4D, middle). We also found significant inverse correlations between K/T and subset 1/Treg and subset 2/Treg (Figure 4D & E). Importantly, we found no correlation between K/T and Th1*, Th1, or IL17^+^ cell frequencies (Fig. S5A & B and data not shown). Although we confirmed high IDO activity (Figure 4) and found a reduction in circulating Th17 (especially subset 1) cells in HIV-ART compared to HIV uninfected people (Figure 2), Tregs and the Th17/Treg ratio were similar in the two groups (Figure S3). Thus, increased IDO activity is associated with a reduction in circulating Th17 cell subsets, but the frequency of IL17-producing CD4 T cells does not correlate with IDO activity.

## Discussion

The protective role of IL17-producing CD4 T cells against *Mtb* infection has been demonstrated across species; mice [60–62], non-human primates [63–65], and humans[1–3]. Studies that demonstrate Th17 cells contribute to the control of *Mtb* infection in humans used different approaches to identify the cells. We used a combination of cell markers [12,15–17,20,21] to broadly characterize Th17 cells and then confirmed that they produce IL17, instead of using IL17 production alone to define Th17 cells [3,66]. Since there are cytokine-independent mechanisms of T cell responses, and since cytokine production can be influenced by stimulation conditions, our approach affords a broader characterization of Th17 cells to determine the impact of HIV coinfection. We also considered other available techniques for studying Th17 cells such as sequencing approaches. Our approach of first identifying Th17 cells based on cell surface markers is compatible with single-cell sequencing that has revolutionized the study of CD4 T cells beyond cytokine production. The existing technologies do not reliably allow the sequencing of cells after fixation and permeabilization for the detection of cytokines.

A recent study in macaques showed granulomas that controlled *Mtb* were enriched for T1T17 (Th1*) cells that did not express IL17 transcripts [65]. Furthermore, since *Mtb*-antigen specific cells must traffic to sites of disease, using chemokine receptor expression alone in blood may underreport the population of these cells. Our finding of disproportionate reduction of specific circulating Th17 cell categories in HIV-ART reveals the importance of using multiple criteria to identify CD4 T cells of interest, and that ART-mediated suppression of HIV does not reconstitute all Th17 cell types equally. Previous reports indicate that CD4^+^CD161^+^ T cells represent a subset of fast acting T cells that inhibit mycobacterial growth in unexposed humans but not TB patients [67] and are also significantly reduced in active TB compared to latent TB [68], suggesting their protective immunity against *Mtb* infection. Although these studies did not include CD26, they may probably constitute subset 1 Th17 cells identified in our study. The preferential production of IL17 by Th17 subsets 1 and 2 depending on the type of stimulation reiterates the need to study pathogen-specific responses using the antigens specific to the pathogen of interest, and that responses to polyclonal stimulation with superantigens may overestimate responses by distinct T cell subsets. Studies to identify CD4 T cell defects that contribute to TB susceptibility in HIV and HIV-ART will benefit from higher resolution characterization of CD4 T cell subsets.

Our finding of a higher magnitude of IL17-than IFNγ-producing CD4^+^ T cells in HIV-ART upon Mtb300 stimulation is contrary to the impression that IFNγ production is the dominant response to *Mtb* infection [69] but agrees with a recent study that revealed evidence that people who have been exposed to *Mtb* but remain negative by IFNγ responses have detectable responses to *Mtb* antigens when other readouts are used [70]. Until recently, those individuals were thought to remain uninfected by *Mtb*; our results add weight to the evolving concept that considering IFNγ as the canonical response to *Mtb* infection requires reconsideration.

Although Mtb300 peptide megapool we used in our study contains peptides from multiple *Mtb* antigens including ESAT6 and CFP10, the sensitivity of our assay may be lower compared to the IGRA test because QuantiFERON-TB Gold Plus blood test is designed to detect both CD4 and CD8 T cells and uses whole blood; we only report on CD4 T cells in PBMCs. Our findings indicate that IL17 responses to *Mtb* but not CMV antigens or SEB can be preserved in HIV-ART. We hypothesize that IL17-producing cells may be less activated than IFNγ-producing cells and/or express the CCR5 coreceptor at lower levels or at lower frequencies and are thus preserved from HIV infection. The technical challenge of optimal measurement of T cell activation and cytokine production in one condition (detection of cytokines requires protein transport inhibitors which hinder T cell activation marker expression) precluded us from testing this hypothesis.

The K/T ratio is a strong predictor of mortality risk in ART-suppressed PWH [71], does not normalize in PWH initiating ART early, and durably suppressed early-ART initiators have high plasma KT ratios compared with HIV-negative controls [31]. Therefore, the elevated tryptophan catabolism in HIV-ART reported here and previously [27,31] may explain, in part, the persistently increased risk of tuberculosis in treated HIV. Inhibition of IDO-1 with 1-methyl-D-tryptophan (indoximod) in macaques was reported to decrease *Mtb* burdens and pathology, and increased the proliferation of CD4^+^ and CD8^+^ memory T cells [72] Together, these results suggest that enhancing Th17 cell responses by IDO inhibition may be beneficial in the context of TB.

Our study has limitations: 1) the participants were from a study investigating the long-term clinical and immunological consequences of HIV infections and their treatment, therefore, we had few HIV uninfected participants; 2) 10 of 13 PWH initially with a negative TST prior to TST conversion had high viral loads that probably reduced the sensitivity of diagnosis; 3) INH treatment was not given as directly observed therapy (DOT) and we could not ascertain treatment compliance by the study participants. Nevertheless, we found that Th17 cells are a heterogenous population and IL17 responses to *Mtb* are preserved in HIV-TB coinfected individuals, and we demonstrated the link between IDO activity and circulating *Mtb*-responsive Th17 cells in humans. With IDO-1 inhibitors in clinical trials [73], management of TB disease might benefit from simultaneous therapeutic inhibition of IDO-1 activity to enhance Th17 cell responses, particularly in PWH where HIV further enhances tryptophan catabolism via the kynurenine pathway.

## 3 Author Contributions

PO: Conceptualization, Data curation, Formal analysis, Investigation, Methodology, Writing – original draft, review & editing, Funding acquisition. AT: Data Curation, Methodology. FM: Data Curation, Methodology. DG: Data Curation, Methodology. MK: Data Curation, Resources. FA: Data Curation, Methodology. CSLA: Resources. JNM: Resources. SGD: Resources. PWH: Conceptualization, Supervision, Resources, Writing – review & editing, Funding acquisition. JDE: Conceptualization, Supervision, Resources, Writing – review & editing, Funding acquisition.

All authors approved the final version of the manuscript.

## Acknowledgments

We thank the study participants in the SCOPE study for providing samples, UCSF Core Immunology Lab and UCSF-CFAR Specimen Processing & Banking Subcore.

