## Supplementary material for "High-parameter phenotypic characterization reveals a subset of human Th17 cells that preferentially produce IL17 against *M. tuberculosis* antigen"

### Supplementary Figures

**Supplementary Figure 1:**

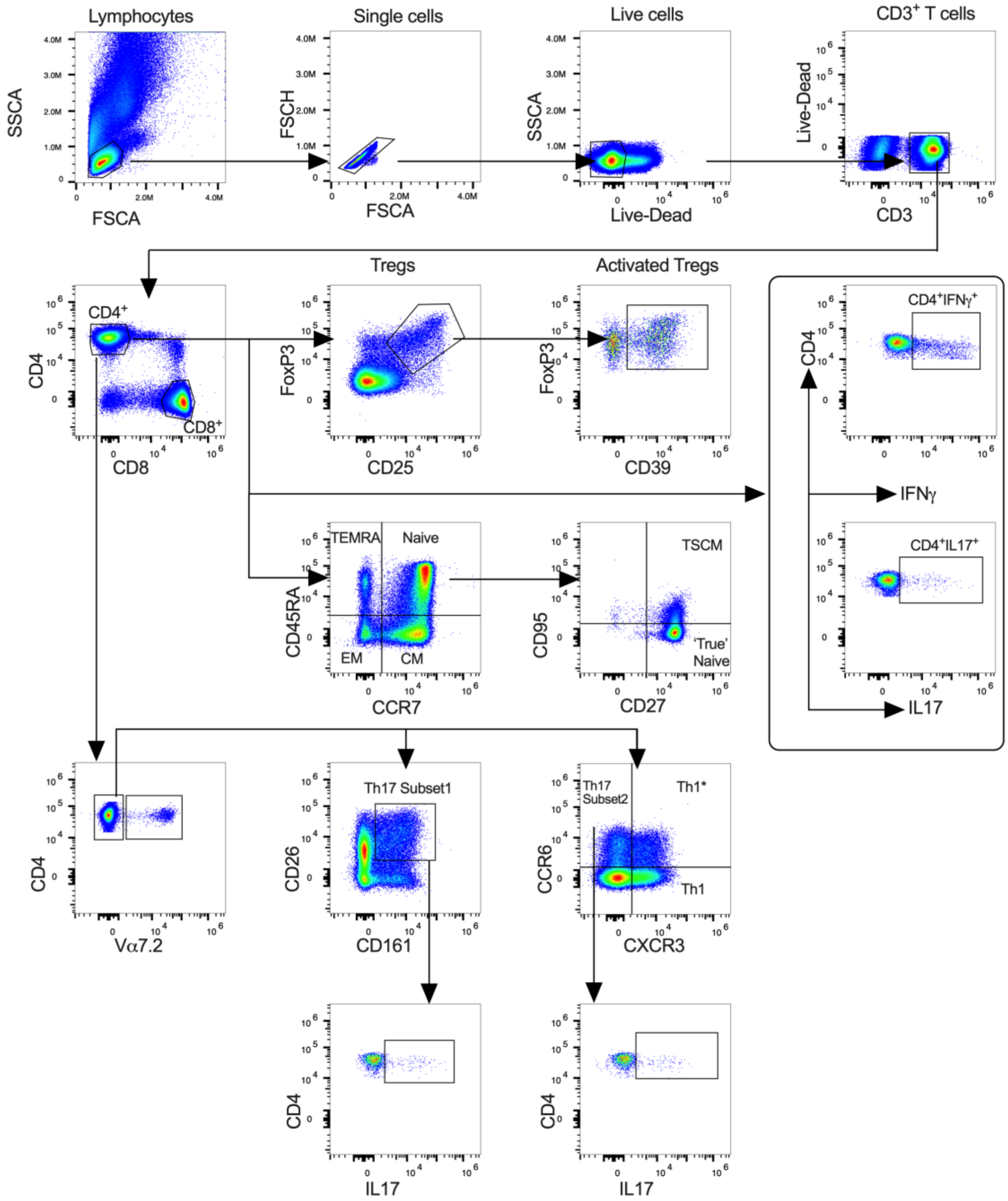


**Supplementary figure 1: Gating strategy to identify Th17 cells subsets and characterize Mtb-antigen specific T cells.** Th17 subset 1 defined as CD4$^+$V$\alpha$7.2$^-$CD26$^+$CD161$^+$; Th17 subset 2 is defined as CD4$^+$V$\alpha$7.2$^-$CCR6$^+$CXCR3$^-$ and Th1* (Th1Th17) defined as CD4$^+$V$\alpha$7.2$^-$CCR6$^+$CXCR3$^+$. T cell differentiation defined based on expression of CD45RA and CCR7 as naive, central memory (CM), effector memory (EM) and terminally differentiated (TEMRA) cells. TSCM = Stem cell like memory cells.

**Supplementary Figure 2:**

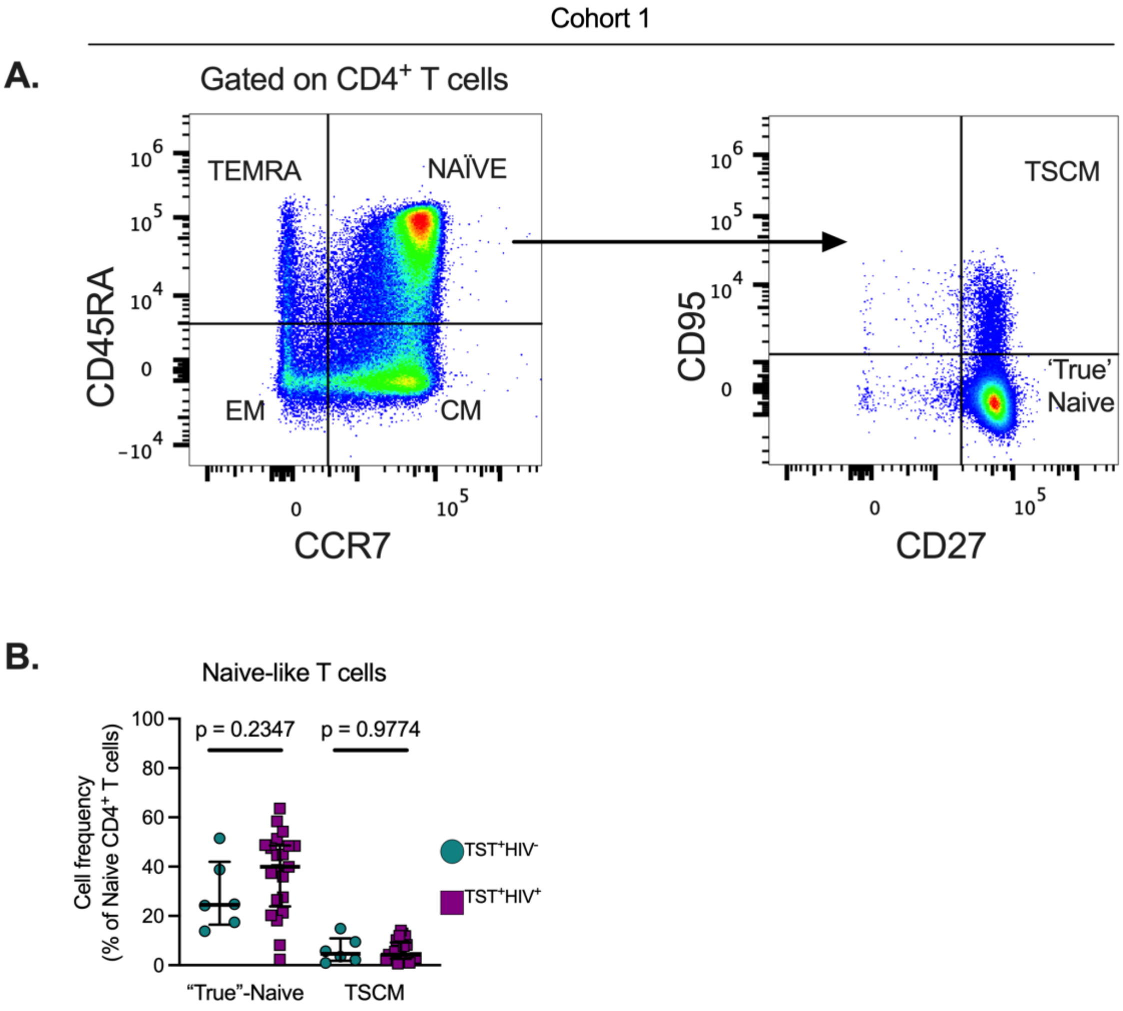


**Supplementary figure 2: Similar circulating stem-cell like memory T cells in PBMC of LTBI individuals with treated HIV or without HIV. (A)** Representative flow cytometry plots for identification of T cell differentiation states; TEMRA (terminally differentiated effector cells), EM (effector memory), CM (central memory), TSCM (Stem Cell like memory). **(B)** Frequency of Naive-like $CD4^+$ T cells gated from $CD3^+CD4^+$ T cells. Statistics: Mann-Whitney test.

**Supplementary Figure 3:**

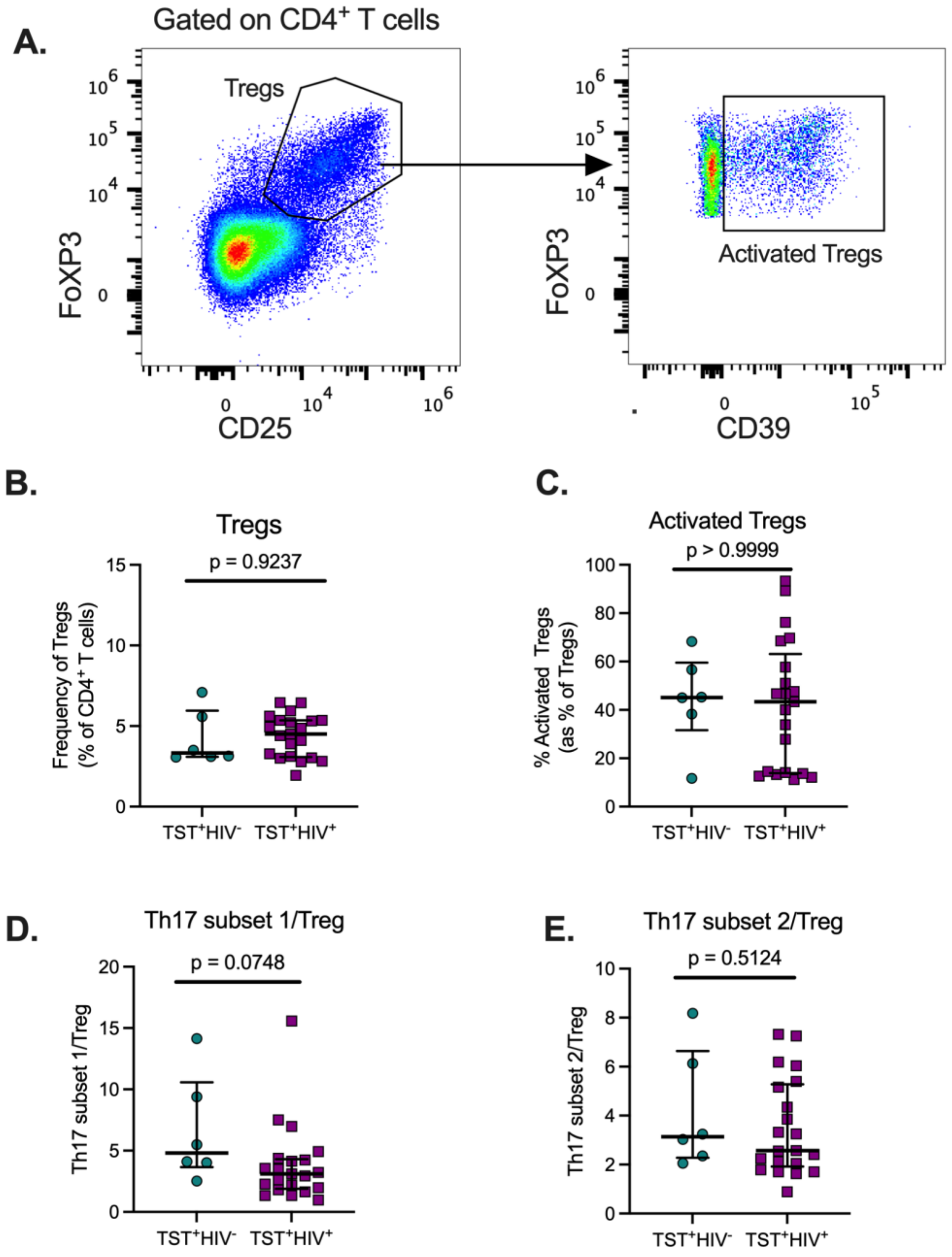


**Supplementary figure 3: Frequency of circulating Tregs and Th17/ Treg in LTBI individuals with treated HIV or without HIV. (A)** Representative flow cytometry plots for identification of Tregs and activated Tregs. **(B)** Frequency of Treg cells gated on $CD4^+$ T cells. **(C)** Activated Treg cells as a fraction of total Tregs. **(D)** subset 1/Treg ratio when T helper 17 cells are identified based on co-expression of CD26 and CD161. **(E)** subset 2/Treg ratio when T helper 17 cells are identified based on expression of chemokine receptor CCR6. Subset 1 and subset 2 Th17 cells are pre-gated on $CD4^+V\alpha7.2^-$ cells. Statistics: Mann-Whitney test.

### Supplementary Figure 4:

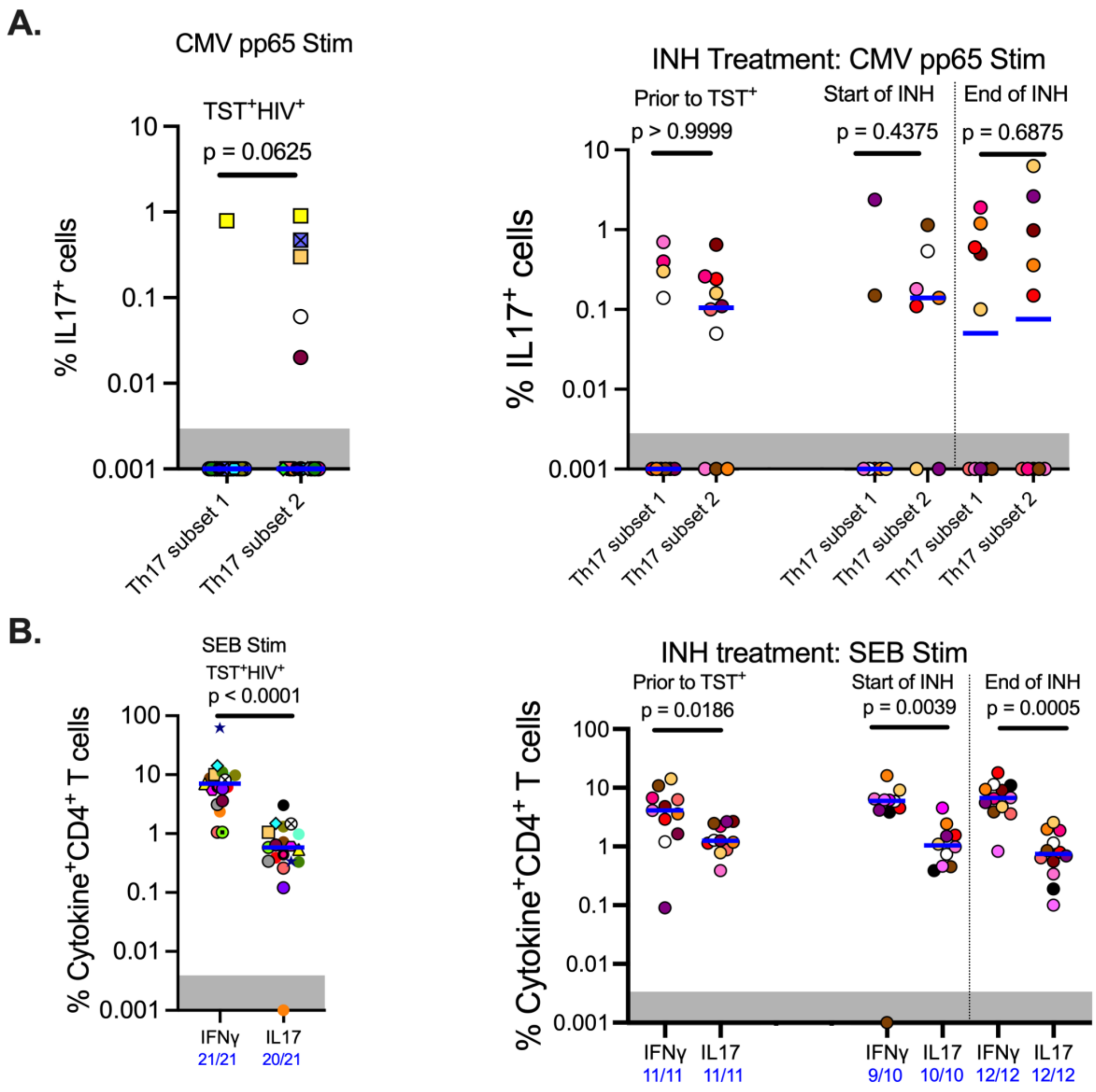


**Supplementary figure 4: CMV-responsive IL17 producing cells are enriched in subser 2 compared with subset 1 cells and higher IFNg than IL17 production to SEB.** PBMC were stimulated with CMV pp65 peptide pool (**A**), or (SEB) (**B**) and cytokine production measured by intracellular cytokine staining. IL17 producing subset 2 and subset 1 Th17 cells to CMV pp65 (**A**). IL17 or IFNγ producing CD4$^{+}$ T cells (bulk response) to SEB (**B**); left panels = LTBI infected individuals with treated HIV, right panels = longitudinal analysis of HIV infected individuals on ART (with varying plasma viral load) who were TST negative then converted to TST positive sampled at the start and end of isoniazid treatment. Values are after subtraction of background response from no stimulation wells; values within grey area depict undetectable responses above the background. Fractions on the x-axis indicate participants with detectable cytokine response. The blue lines indicate median response. Statistics - Wilcoxon test (paired) or Mann-Whitney test (unpaired).

**Supplementary Figure 5:**

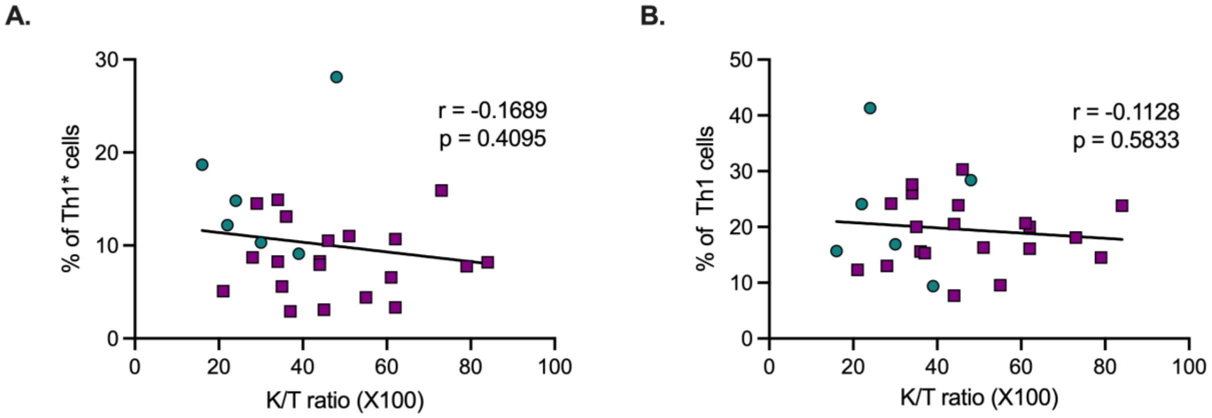


**Supplementary figure 5: Association between IDO-1 activity and circulating Th1* and Th1 cells.** Frequency of Th1* cells (**A**) and Th1 cells (**B)** in PBMC measured by spectral flow cytometry and matched plasma Tryptophan (T) and Kynurenine (K) concentration was measured by Mass Spectrometry and IDO-1 activity estimated as K/T ratio. Cross sectional analysis of LTBI infected individuals with treated HIV (plum squares) or without HIV (teal circles). Statistics - Spearman correlation test.

**Supplementary Figure 6:**

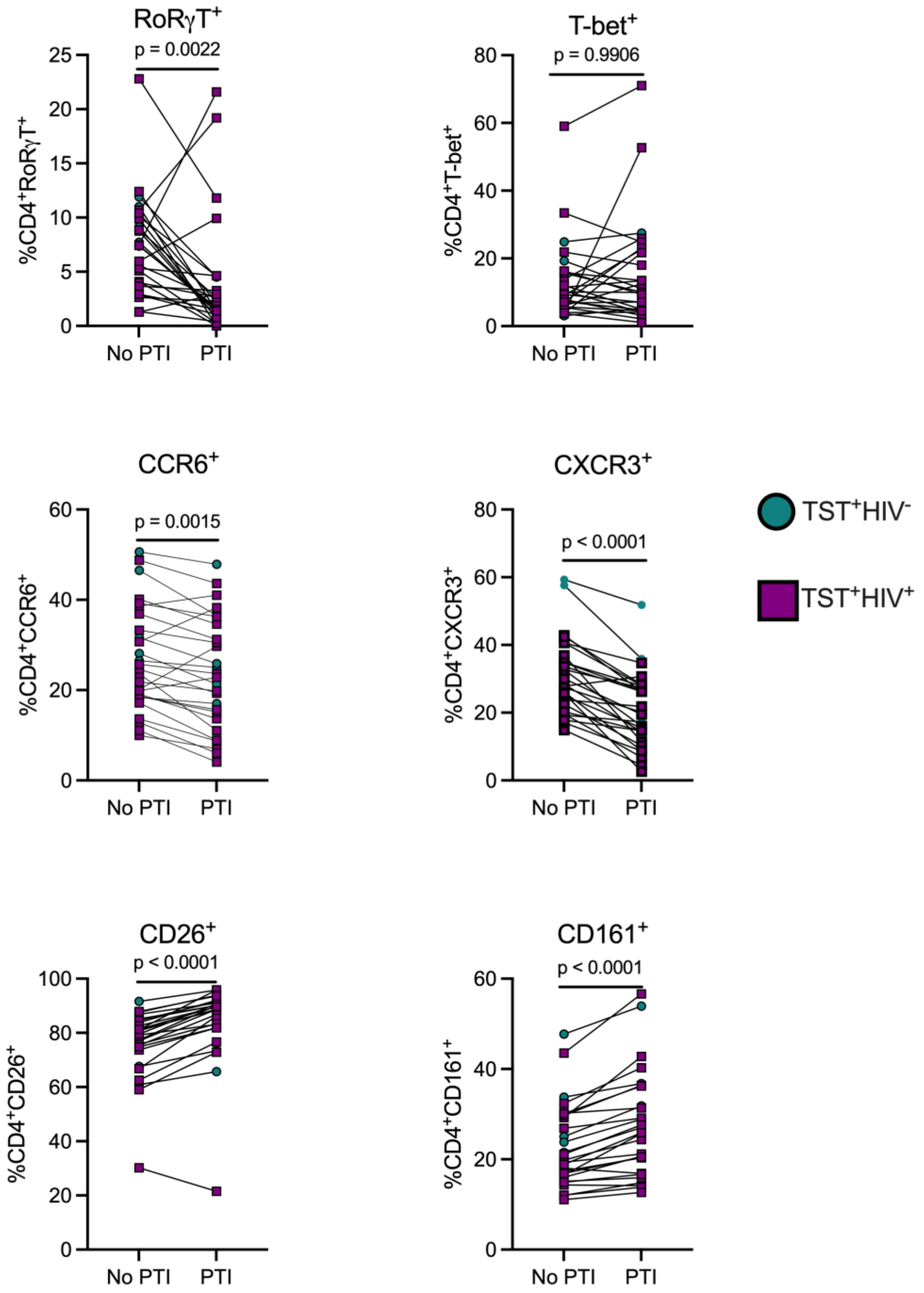


**Supplementary figure 6: The effect of protein transport inhibitors (PTI) on expression of markers of Th17 cells.** Cells were split into two equal portions and cultured for a total of 20 hours in the absence of antigen. In one portion, Golgi stop and Golgi plup (protein transport inhibitors, PTI) was added in the last 18 hours while the portion received no PTI. Cells were stained and analysed by spectral flow cytometry. Statistics: Wilcoxon test.

**Supplementary Figure 7:**

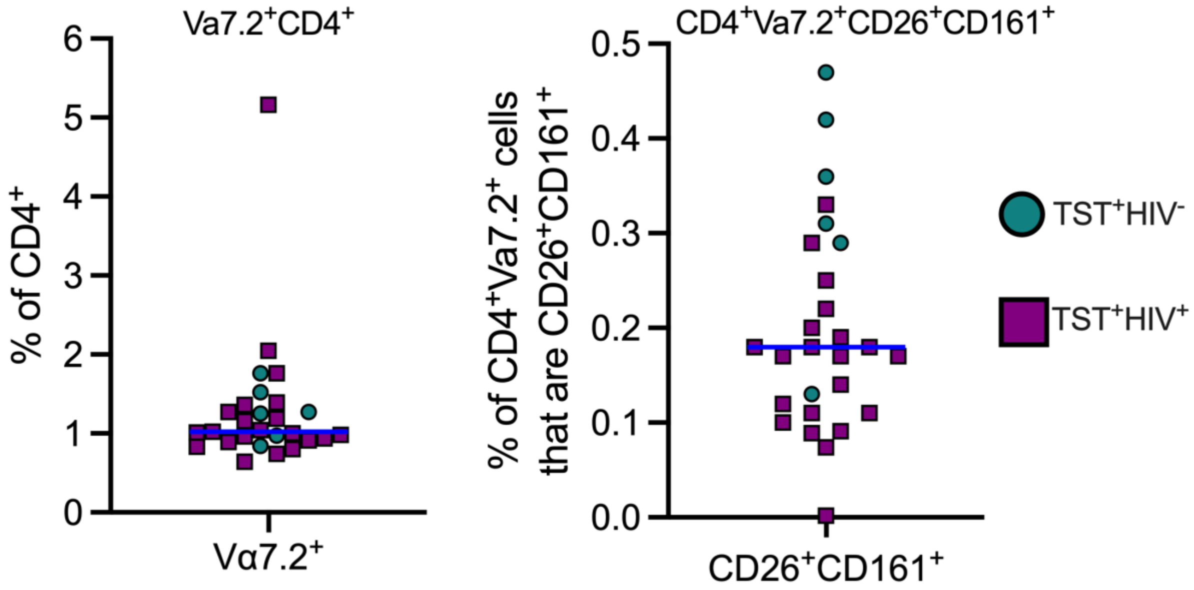


**Supplementary figure 7: Frequency of circulating CD4⁺ mucosa associated invariant T cells (MAIT).** CD4⁺ MAIT cells are defined by canonical expression of T cell antigen receptor (TCR) Vα7.2 (also refered to as TRAV1-2). In addition, MAIT cells express cell surface markers CD161 and CD26. Blue horizontal line shows the median value.
